# Growth and selection of the cyanobacterium *Synechococcus* sp. PCC 7002 using alternative nitrogen and phosphorus sources

**DOI:** 10.1101/603373

**Authors:** Tiago Toscano Selão, Artur Włodarczyk, Peter J. Nixon, Birgitta Norling

## Abstract

Cyanobacteria, such as *Synechococcus* sp. PCC 7002 (Syn7002), are promising chassis strains for “green” biotechnological applications as they can be grown in seawater using oxygenic photosynthesis to fix carbon dioxide into biomass. Their other major nutritional requirements for efficient growth are sources of nitrogen (N) and phosphorus (P). As these organisms are more economically cultivated in outdoor open systems, there is a need to develop cost-effective approaches to prevent the growth of contaminating organisms, especially as the use of antibiotic selection markers is neither economically feasible nor ecologically desirable due to the risk of horizontal gene transfer. Here we have introduced a synthetic melamine degradation pathway into Syn7002 and evolved the resulting strain to efficiently use the nitrogen-rich xenobiotic compound melamine as the sole N source. We also show that expression of phosphite dehydrogenase in the absence of its cognate phosphite transporter permits growth of Syn7002 on phosphite and can be used as a selectable marker in Syn7002. We combined these two strategies to generate a strain that can grow on melamine and phosphite as sole N and P sources, respectively. This strain is able to resist deliberate contamination in large excess and should be a useful chassis for metabolic engineering and biotechnological applications using cyanobacteria.

## 1. Introduction

As part of the effort to advance a carbon-neutral economy, cyanobacteria are increasingly being used for metabolic engineering. They are photoautotrophic organisms, able to grow with rather simple requirements – minimal media with inorganic nitrogen and phosphorus sources, using light for energy generation and CO_2_ as the sole carbon input – and, in recent years, have been shown to be able to produce a plethora of different molecules, from commodity chemicals such as lactate or ethanol (Angermayr et al., 2012; Dexter et al., 2015; Gordon et al., 2016), to biofuels (e.g., free fatty acids (Kato et al., 2017; Ruffing, 2014) or butanol (Fathima et al., 2018; Shabestary et al., 2018)) to speciality chemicals such as farnesene (Halfmann et al., 2014), squalene (Choi et al., 2017; Englund et al., 2014) or limonene (Wang et al., 2016). Several strains are commonly used in both basic as well as applied research, with fast-growing marine strains such as *Synechococcus* sp. PCC 7002 (henceforth “Syn7002”) being of special interest. As it is able to grow in seawater (thus not competing for freshwater resources), withstand high light intensities and temperatures up to 40 °C (useful in large scale open-air facilities), is naturally transformable, has an optimal division time of roughly 4 hours, an available genome sequence as well as several synthetic biology and genetic modification tools (Begemann et al., 2013; Clark et al., 2018; Frigaard et al., 2004; Gordon et al., 2016; Ludwig and Bryant, 2011; Ludwig and Bryant, 2012; Markley et al., 2015; Perez et al., 2016; Xu et al., 2011), this strain is ideally poised to become an important biotechnological workhorse.

Large scale cyanobacterial cultivation can be done in either semi-enclosed systems, such as hanging bags, or in open systems, be it raceway ponds or air-lifted stock ponds (Schoepp et al., 2014). Closed systems have the advantage of higher controllability, lesser chance for culture contamination and generally higher growth yields. However, they are substantially costlier to operate than open systems, with either air-lifted stock ponds or raceway ponds the most economically viable alternatives found so far (Schoepp et al., 2014). Open systems have, on the other hand, the obvious disadvantage of being exposed to the environment and, as such, are much more prone to contamination. Operating a large open system with cyanobacterial strains carrying antibiotic-resistance genes would also introduce a severe biohazard should the cultures escape and, through horizontal gene transfer, contribute to the spreading of antibiotic resistance to environmental pathogenic species.

Recently, the use of ecologically rare or xenobiotic sources of macronutrients has been explored as a means to generate selective pressure towards the growth of genetically modified organisms without the use of antibiotics (Kanda et al., 2014; Loera-Quezada et al., 2016; Pandeya et al., 2017; Polyviou et al., 2015; Shaw et al., 2016) as well as to allow genetically modified plants to outcompete weeds, while consuming considerably less phosphorus (Lopez-Arredondo and Herrera-Estrella, 2012). To this end, phosphite dehydrogenase (PtxD), an enzyme converting phosphite, an ecologically rare form of phosphorus, into phosphate, has been introduced into a variety of organisms, and in some cases used as a selectable marker (Kanda et al., 2014; Lopez-Arredondo and Herrera-Estrella, 2012; Nahampun et al., 2016; Pandeya et al., 2017). A synthetic pathway to utilize melamine, a xenobiotic nitrogen-rich compound, has also been devised and introduced into different organisms (Shaw et al., 2016). Introduction of the complete pathway (consisting of 6 enzymes) in *Escherichia coli* allowed the carrying strain to overcome deliberate contamination (Shaw et al., 2016).

In many cases, these pathways or genes were introduced into target organisms with the help of antibiotic cassettes (Loera-Quezada et al., 2016; Motomura et al., 2018; Shaw et al., 2016) and even though this proves that the pathways confer an advantage to the organisms carrying them, the risk of horizontal gene transfer of the antibiotic resistance cassette(s) still exists. With that in mind, we set out to develop strains of Syn7002 that would be able to grow on melamine and phosphite as sole N and Pi sources but using metabolic selection to drive their genomic integration. We were able to obtain, through laboratory evolution, six different Syn7002 strains that can grow on melamine as a sole N source and validated the use of the *ptxD* gene and phosphite as an efficient selectable marker in this cyanobacterial species. We further engineered one of the best melamine-growing strains by introducing the *ptxD* gene and show that this strain, which is able to grow using both melamine and phosphite as N and Pi sources, respectively, is able to withstand and easily outcompete deliberate contamination, even in large excess, thus paving the way for economically feasible and antibiotic-less outdoor large scale growth of these modified photosynthetic microorganisms.

## 2. Material and Methods

### 2.1 Cell growth conditions

*Synechococcus* sp. PCC 7002 (a kind gift from Prof. Donald Bryant, Penn State University, USA) was grown photoautotrophically in medium A (Stevens et al., 1973) using D7 micronutrients (Arnon et al., 1974), supplemented with either 12 mM sodium nitrate (AD7-NO_3_), 4 mM cyanurate (AD7-Cya) or 2 mM melamine (AD7-Mel) as indicated and vitamin B_12_ (0.01 mg.L^−1^). For phosphite-utilizing strains, potassium dihydrogen phosphate (Pho) was substituted by potassium dihydrogen phosphite (Phi, Rudong Huayun Chemical Co., Ltd., Jiangsu, China), at 0.370 mM (Pho 1x or Phi 1x). Solid medium was prepared by supplementing the above media with 1.2% (w/v) Bacto-Agar (BD Diagnostics) and 1 g/L sodium thiosulfate.

For growth experiments, liquid pre-cultures of either Syn7002 WT (grown in AD7-NO_3_) or melamine-growing strains (grown in AD7-Mel) were cultivated at a low-light intensity of 50 µmol photons·m^−2^·s^−1^ at 38 °C, 1% CO_2_, 160 rpm, until an OD_730_ between 4 and 6 (late logarithmic phase under low-light conditions). Cells were pelleted and washed twice with AD7-Mel medium without phosphate (AD7-Mel P-) prior to inoculation in baffled flasks. Liquid cultures (25 mL total volume, three biological replicates per strain) were grown in 100 mL baffled Erlenmeyer flasks, in a 740-FHC LED incubator (HiPoint Corporation, Taiwan), at 38 °C in air supplemented with 1% (v/v) CO_2_, under constant illumination of 300 µmol photons·m^−2^·s^−1^, using an LED Z4 panel, set to 215 µmol photons·m^−2^·s^−1^ of red light (660 nm), 50 µmol photons·m^−2^·s^−1^ of green light (520 nm) and 35 µmol photons·m^−2^·s^−1^ of blue light (450 nm) and shaken at 200 rpm. Cell growth was monitored by measuring the optical density at 730 nm (OD_730_) in a 1-cm light path with a Cary 300Bio (Varian) spectrophotometer. For dry cell weight determination, 1 to 2 mL culture volume at a determined OD_730_ (between 8 and 10) were filtered onto pre-dried and pre-weighed glass microfiber filters (47 mm diameter, 1 µm pore size, GE Healthcare, Cat. No. 1822-047), washed twice with deionized water and dried overnight at 65 °C (until mass deviation between readings was no higher than 0.0001 g). All measurements were performed using biological triplicates of each strain.

Supercompetent *Escherichia coli* cells (Stellar, TaKaRa) were used for construction of all relevant plasmids and were cultured in LB medium, at 37 °C, supplemented with either 50 µg·mL^−1^ kanamycin, 50 µg·mL^−1^ spectinomycin, 50 µg·mL^−1^ gentamycin or 100 µg·mL^−1^ carbenicillin, as appropriate. Unless otherwise specified all chemicals utilized were procured from Sigma-Aldrich.

### 2.2. Strain construction

#### 2.2.1. Melamine growing strain

All PCR reactions were performed using Q5 DNA polymerase (New England Biolabs, NEB) unless otherwise specified and PCR products were routinely digested overnight with DpnI and purified using a the EZ-10 Spin Column PCR Products Purification Kit (BioBasic) prior to DNA assembly. A DNA fragment comprising the *glpK* neutral genomic integration site (Begemann et al., 2013) flanked by 500-bp upstream and downstream regions was PCR amplified from Syn7002 genomic DNA (gDNA) using primers D08807 and D08808 (see Supplemental Table 1). The purified PCR product was ligated into pCR-Blunt II TOPO (Invitrogen) following the manufacturer’s instructions and transformed into chemically competent Stellar *E. coli* cells, resulting in plasmid pCRBlunt-glpK (correct assembly was confirmed by Sanger sequencing using universal M13 primers). Using primers D98496993 and D77036, the pCRBlunt-glpK backbone was reverse-PCR amplified, and the melamine operon was amplified in two halves from a synthetic construct (GenScript, Hong Kong, Ltd.) using primers D98847023 and D98847024 (first half) and D98847025 and D98847026 (second half). Both fragments were assembled into pCRBlunt-glpK using the NEBuilder HiFi DNA Assembly Master Mix (NEB), as per the manufacturer’s instructions. 1 µL of the assembly mix was transformed into Stellar *E. coli* super-competent cells, resulting in plasmid pSJ051. Correct assembly of the melamine operon was confirmed by Sanger sequencing using the primers indicated in Supplemental Table 1. Syn7002 WT was transformed by double homologous recombination as previously described (Frigaard et al., 2004), with modifications. Briefly, 2 µg of pSJ051 were used to transform 2 mL of a Syn7002 culture at an OD_730_ of 0.5 and incubated overnight, as described above, in a 12 mL round bottom snap-cap tube. The following day the culture was spun down, the pellet resuspended in 50 µL of the supernatant prior to spreading on an AD7-Cya plate, to favour integration of the entire melamine operon into the *glpK* site. This plate was incubated under the conditions described above until colonies appeared (after 2 weeks). Eight colonies were picked and restreaked 4 times on AD7-Cya plates and, subsequently, on AD7-Mel plates until full chromosomal segregation, tested using primers D99280067 and D99280068 (see Figure 1), was confirmed. Only six of the initial eight colonies survived, having evolved into strains Mel1, Mel4, Mel5, Mel6, Mel7 and Mel8.

**Figure 1.**
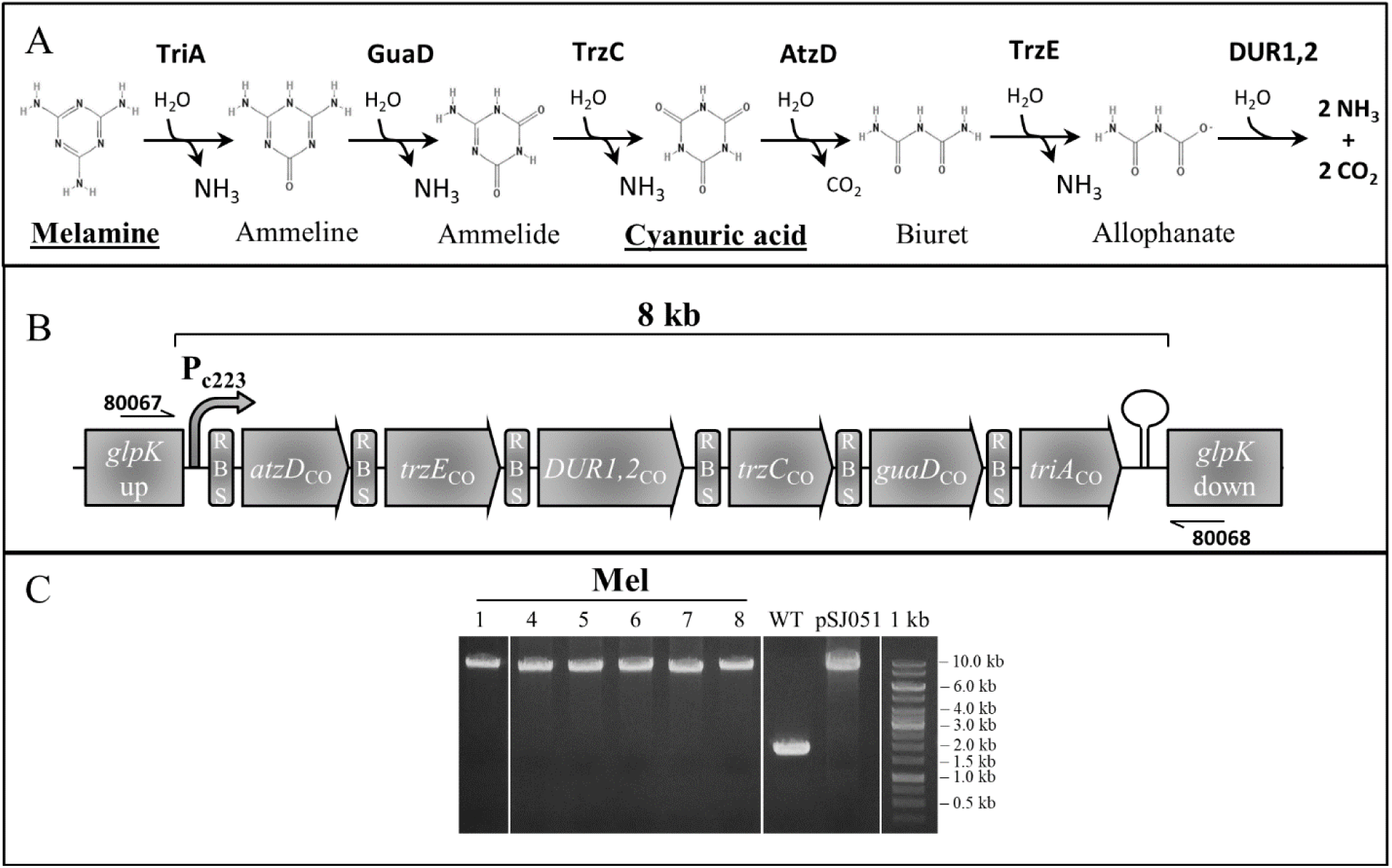
Overview of melamine selection tool. **A.** Melamine utilization pathway reactions. One mol of melamine yields 6 mol ammonia and 3 mol carbon dioxide. **B.** Schematic view of the melamine utilization operon. Primers indicated were used to confirm full genome integration of the pathway. Different parts are not to scale **C.** 0.6% agarose gel of PCR reaction using primers stated in A. (see Supplemental Table 1 for sequences)

#### 2.2.2. Phosphite–utilizing and combined melamine/phosphite-utilizing strains

A region 500 bp upstream to 500 bp downstream of the neutral genomic integration site between ORF *A0935* and *A0936* (Davies et al., 2014) was PCR amplified from Syn7002 gDNA using primers D100023580 and D100023581. pUC19 (Invitrogen) was digested with XbaI (NEB) and purified from the agarose gel band using the EZ-10 Spin Column DNA Gel Extraction Kit (BioBasic). The *A0935-A0936* site was assembled into the digested pUC19 backbone using the pEASY-Uni Seamless Cloning and Assembly Kit (TransGen Biotech Co., Ltd, China), according to the manufacturer’s instructions, and transformed into Stellar *E. coli* cells, resulting in plasmid pSZT001. Primer pair A0935_UCO_F and A0936_UCO_R were used to reverse-PCR amplify the pSZT001 backbone and pair D98496996 and D100141467 to amplify a synthetic, codon-optimized version (by GenScript, Hong Kong, Ltd) of the *Pseudomonas stutzeri* WM88 phosphite dehydrogenase (*ptxD*) gene (Loera-Quezada et al., 2016), driven by the *Amaranthus hybridus* constitutive *psbA* promoter (Elhai and Wolk, 1988). Both fragments were assembled using the pEASY-Uni kit, as described above, resulting in plasmid pSJ135. 2 µg of this plasmid were used to transform both Syn7002 WT as well as the Mel5 strain, as described above, with the exception that, prior to transformation, cultures were spun down and washed twice with AD7 medium lacking phosphate (AD7-NO_3_ P-for WT and AD7-Mel P-for Mel5) and subsequently plated onto either AD7-NO_3_ Phi 1x (0.370 mM phosphite) or AD7-Mel Phi 1x, respectively. Both plates were incubated under the conditions described above until colonies appeared (10 days). Eight colonies from each plate were picked and consecutively restreaked on AD7-NO_3_ Phi 1x or AD7-Mel Phi 1x until full chromosomal segregation was confirmed by colony PCR using primers D100043610 and D100043611 (see Fig. 4), resulting in strains A0935-ptxD and Mel5-ptxD, respectively.

To further test the use of phosphite as a selectable marker, a DNA fragment containing an YFP gene under the control of the strong constitutive P_cpt_ promoter (Markley et al., 2015) was amplified from pAcsA-cpt-YFP (a kind gift from Prof. Brian Pfleger, University of Wisconsin-Madison, USA) using primers D100263687 and D100263688. The pSJ135 backbone was reverse-PCR amplified using primers D100141467 and D100098818 and the two fragments assembled using the pEASY-Uni kit, yielding pSJ141. Transformation of either Syn7002 WT or Mel5, using phosphite media for selection of transformants, and selection of fully segregated strains by colony PCR was performed as described above.

### 2.3. Knock-out of putative phosphonate transporters

Putative phosphonate transporter genes were identified by using the BlastP tool within CyanoBase (http://genome.microbedb.jp/blast/blast_search/cyanobase/genes), limiting the search to Syn7002 and using as a search template the amino acid sequence of PhnD from *Prochlorococcus marinus* sp. MIT9301 (Bisson et al., 2017; Feingersch et al., 2012). Two homologues (annotated as putative phosphate/phosphonate-binding ABC transporters) were identified in Syn7002: A0336 (E value = 1e^−96^) and G0143 (E value = 2e^−08^). DNA sequences 500 up-and downstream of these genes were amplified from Syn7002 cells using primers D15106 and D15107 (A0336 upstream region), D15108 and D15109 (A0336 downstream region), D15110 and D15111 (G0143 upstream region) and D15112 and D15113 (G0143 downstream region). Purified DNA fragments were assembled into an XbaI-digested pUC19 fragment with either a spectinomycin-resistance cassette (in the case of *A0336*), amplified from pBAD42 (using primers D98646038 and D98646039) or a gentamycin-resistance cassette (in the case of *G0143*), amplified from pVZ322 (using primers D99047654 and D99047655), using the NEBuilder HiFi DNA Assembly Master Mix, according to the manufacturer’s instructions, resulting in pSJ156 (pUC19-Δ*A0336*::SpR) and pSJ157 (pUC19-Δ*G0143*::GmR). Both plasmids were used to transform Syn7002 WT and the A0935-ptxD strain and, upon full segregation, pSJ157 was used to transform Δ*A0336*::SpR deletion strains in the WT (WTΔ*A0336* strain) and A0935-ptxD (ptxDΔ*A0336* strain) backgrounds, resulting in double-knockout strains WTΔ*A0336*Δ*G0143* and ptxDΔ*A0336*ΔG*0143*. WT Syn7002, A0935-ptxD, ptxDΔ*A0336*, ptxDΔ*D0143* (Δ*G0143*::GmR in an A0935-ptxD background) and ptxDΔA*0336*ΔG*0143* were grown in regular AD7 (Pho 1x) medium, in the presence of appropriate antibiotic concentrations (50 µg·mL^−1^ spectinomycin and/or 50 µg·mL^−1^ gentamycin), under the conditions described above. Cultures of all strains were washed twice with AD7-NO_3_ P-, resuspended to an OD_730_ = 4 in the same medium and serially diluted 1:10 (from 4×10^0^ to 4×10^−5^) using the same AD7-NO_3_ P-medium. 10 µL of each dilution were spotted on either AD7-NO_3_ Pho 1x or AD7-NO_3_ Phi 20x, cultured under the conditions described above, for 5 days.

### 2.4. YFP fluorescence measurement

Whole cell YFP fluorescence was determined for triplicate cultures (15 mL each) grown in regular AD7 medium to an OD_730_ between 0.5 and 1, with 150 µL aliquots measured using 96-well black clear bottom plates, in a Hidex Sense (excitation: 485/10 nm; emission: 535/20 nm). Fluorescence was measured in triplicates for each culture and normalized to OD_730_ (measured at the same time by the Hidex Sense plate reader, as described above), using AD7 medium as blank control.

### 2.5. Genome sequencing

Genomic DNA was prepared from both the WT strain as well as the different melamine-utilizing strains by using the Quick-DNA Fungal/Bacterial Kit (Zymo Research). Library preparation was performed according to Illumina’s TruSeq Nano DNA Sample Preparation protocol. The samples were sheared on a Covaris E220 to ∼550bp, following the manufacturer’s recommendation, and uniquely tagged with one of Illumina’s TruSeq LT DNA barcodes to enable sample pooling for sequencing. Finished libraries were quantitated using Promega’s QuantiFluor dsDNA assay and the average library size was determined on an Agilent Tapestation 4200. Library concentrations were then normalized to 4nM and validated by qPCR on a QuantStudio-3 real-time PCR system (Applied Biosystems), using the Kapa library quantification kit for Illumina platforms (Kapa Biosystems). The libraries were then pooled at equimolar concentrations and sequenced on the Illumina MiSeq platform at a read-length of 300bp paired-end. Genomes were assembled and compared using the Geneious 11.1.4 software (Biomatters Ltd.).

### 2.6. RBS point mutation test

To assess the effect of the RBS change in Mel5, the original pSJ051 plasmid was mutated at the RBS upstream of *triA* (changed from AGGAGA to AGAAGA) by inverse PCR (Liu and Naismith, 2008) using Q5 DNA polymerase (NEB) using primers D101108989, resulting in plasmid pSJ155. This plasmid was used to transform Syn7002 WT, using AD7-Cya and AD7-Mel plates, as described above. The resulting strain, Re-Mel5, was restreaked twice on AD7-Mel plates and tested for growth in AD7-Mel as described above.

### 2.7. Co-culturing competition experiments and flow cytometry

Growth competition experiments were performed by culturing a Syn7002 strain transformed with the pAcsA-cpt-YFP plasmid (constitutively expressing YFP, termed “cptYFP”) and Mel5-ptxD in either AD7-NO_3_ 1xPho or AD7-Mel 20xPhi. Strains were diluted to a starting OD_730_ of 0.05 and their growth (in biological triplicate cultures) followed by flow cytometry using a 3-laser BD LSR Fortessa X20, using the channels of FITC (ext: 488 nm; em: 525/50 nm) to detect YFP-positive cells and APC (ext: 633 nm; em: 670/30 nm) for Chlorophyll a (Chl a)-positive cells. For cell counting cultures were first diluted to an OD_730_ of roughly 0.05 as needed and were acquired on the Fortessa X20 flow cytometer at a consistent rate of 3000 events/s. A log-scale plot of forward scatter (on the x-axis) vs side scatter (on the y-axis) was used as the initial gating to select for live cells and then analysed for Chl a-positive cells. This was then used to draw gates for Chl a-only (Mel5-ptxD) or double positive Chl a/YFP (cptYFP) cells. Cell counts were obtained by acquiring to exhaustion a set sample volume of 50 µL. Cell counts in triplicate samples were derived using the BD FACS Diva Software (v. 8.0) and back-calculated based on the dilution utilized.

### 2.8. Identification of melamine pathway intermediates using LC-MS/MS

Cultures of Syn7002 and melamine-utilizing strains were collected after 48 hours of growth and spun down (14000 g, 5 min, room temperature). Supernatants were filtered through 0.2 µm syringe filters (Acrodisc filters with Supor membrane, PALL) and frozen at -80 °C until further use. Melamine, ammeline, ammelide and cyanuric acid were quantified by LC-MS/MS using a previously described method (Braekevelt et al., 2011) at the NTU Phenomics Centre.

## 3. Results and Discussion

### 3.1. Introduction of melamine degradation pathway requires evolutionary adaptation for efficient usage

The melamine degradation pathway utilized in this study is based on the optimized pathway (including the described R352S mutation in the *guaD* gene product) reported by Shaw and co-workers (Shaw et al., 2016) (Figure 1). In our case we used codon-optimized genes, a synthetic strong cyanobacterial promoter (P_c223_ (Markley et al., 2015)) and the strongest RBS sequence (AGGAGA) tested in Syn7002 (Markley et al., 2015) upstream of all 6 genes. Intergenic regions (21 bp in total), including spacers before (7 bp) and after the RBS sequence (8 bp), were generated by a random DNA sequence generator (http://www.faculty.ucr.edu/~mmaduro/random.htm) and a vector was constructed to target the entire pathway to the *glpK* neutral site (Begemann et al., 2013) of WT Syn7002 (see section 2.2.1 for details). As we wanted to avoid introducing an antibiotic cassette, transformants were selected on plates containing melamine (AD7-Mel plates). However, despite several attempts, it was not possible to obtain colonies when plating directly onto AD7-Mel plates (data not shown). We hypothesised that positive transformants could be selected for by plating cells onto AD7 plates containing cyanuric acid rather than melamine as cyanuric acid is an intermediate in melamine degradation, formed in the 3^rd^ step of the pathway (Fig. 1). This strategy led to the isolation of a small number of cyanuric acid growing colonies. However, these colonies were initially unable to grow on AD7-Mel plates and were therefore restreaked 4 times on AD7-Cya plates before growth could be finally achieved on AD7-Mel plates. Six isolated colonies grown on AD7-Mel plates were analysed further and shown to contain the entire Mel operon and to be fully segregated as judged by PCR analysis (Fig. 1C).

We then compared growth of the different Mel strains and WT Syn 7002 in either AD7-Mel medium or regular AD7 medium (containing nitrate). As can be seen in Fig. 2, the different individual strains, unlike the parental Syn7002 WT strain, were able to grow on melamine as sole nitrogen source, albeit at different rates. Two strains in particular, Mel5 and Mel7, could grow almost as well in AD7-Mel as Syn7002 WT in AD7-NO_3_ while the remaining strains had slower growth and a different colouration (Fig. 2B), a common indication of stress (Collier and Grossman, 1992).

**Figure 2.**
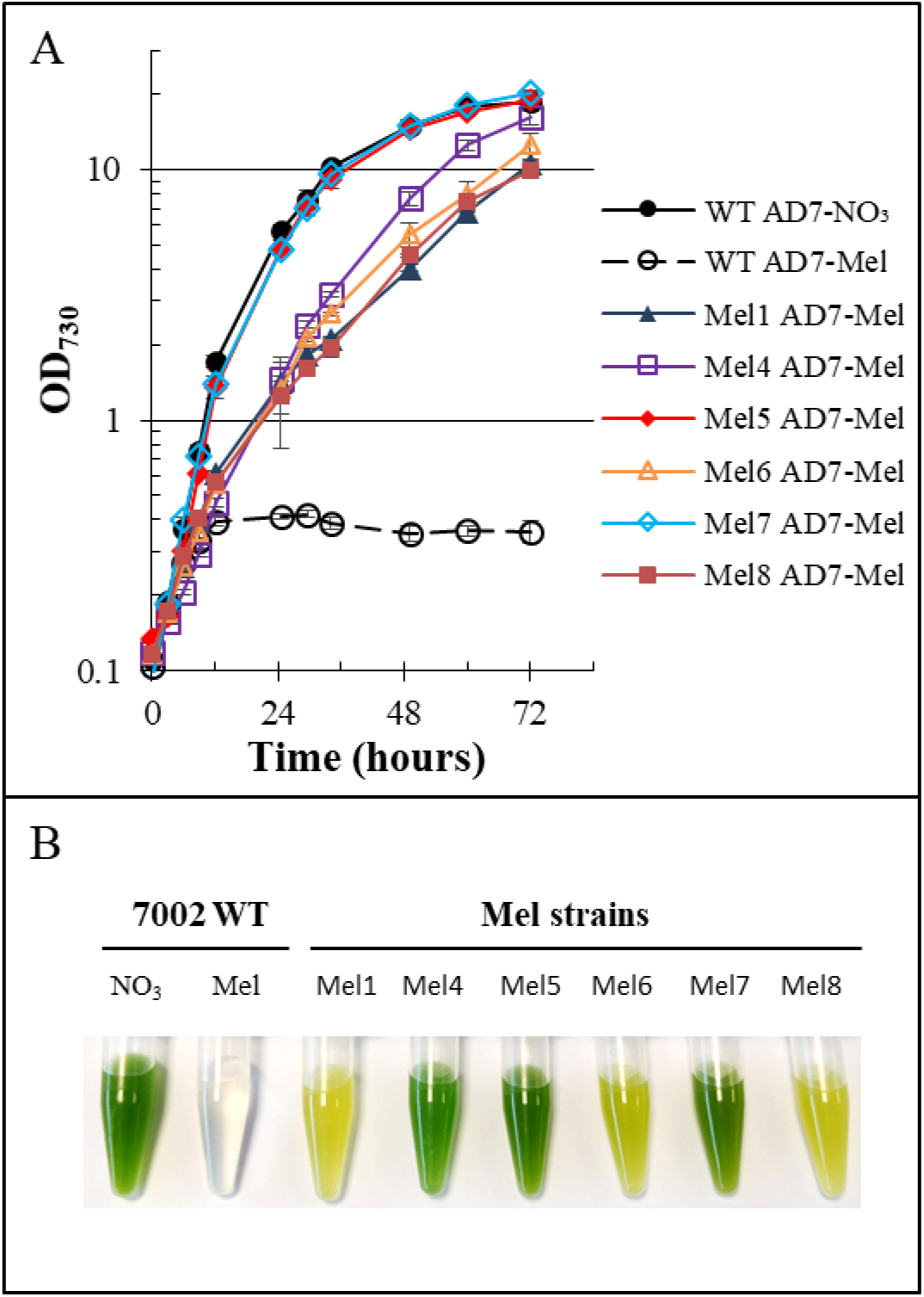
Growth of melamine utilizing strains in melamine containing medium **A**. Growth curve of WT Syn7002 and melamine utilizing strains. **B**. Samples of cultures 48 hours after inoculation. Factors for converting OD_730_ to grams dry cell weight (gDCW)·L^−1^ were calculated for all strains and can be found in Suppl. Table 2.

To further understand the reasons behind these varying phenotypes we sequenced the genomes of all the melamine-growing strains as well as of the Syn7002 WT used in this experiment (obtained from the laboratory of Prof. Donald Bryant, Penn State University). Comparison of the sequences allowed us to pinpoint several mutations in the melamine operon, all of them located either in the RBS preceding the *triA* (encoding melamine deaminase) gene or within the *triA* gene itself (see Supplemental Figure 1). As this is the first step in the melamine degradation pathway, mutations affecting *triA* or its RBS will regulate metabolic flux through the rest of the pathway.

To further clarify the changes occurring in the melamine pathway’s metabolic flux in the different strains, we used LC-MS/MS to quantify the pathway intermediates excreted into the growth medium from melamine to cyanuric acid (see Suppl. Fig. 3). Melamine was very rapidly consumed, within the first 24 hours of growth, in both Mel5 and Mel7, at the same time ammeline (the first intermediate after melamine) was found to accumulate significantly more in Mel5 and Mel7 (86.2±1.6 µM for Mel5 and 57.2±1.5 µM for Mel7) than in the remaining strains (≤25 µM), within the same time frame. Ammelide (the third intermediate) could only be quantified at very low levels (<4 µM detected in Mel5 growth medium after 24 hours) whereas cyanuric acid rapidly accumulated to a concentration of 207.5±16.3 µM in Mel5 and 134.5±8.8 µM in Mel 7 (see Suppl. Fig. 3). The substantial excretion of cyanuric acid into the medium was also reported in the original article on the introduction of the melamine degradation pathway into *E. coli* at levels of 13% of the initial (molar) amount of added melamine (Shaw et al., 2016), strikingly similar to the value of 9.7% observed in the Mel5 strain after 24 hours. It is likely that the mutations present in the remaining strains do not confer as strong an advantage as those found for Mel5 and Mel7, resulting in low intracellular nitrogen levels and poorer growth rates observed under normal light conditions (Fig. 2). It should be noted that pre-cultures, grown under low light, were much less affected, possibly due to a general metabolic rate slowdown under those conditions (data not shown).

**Figure 3.**
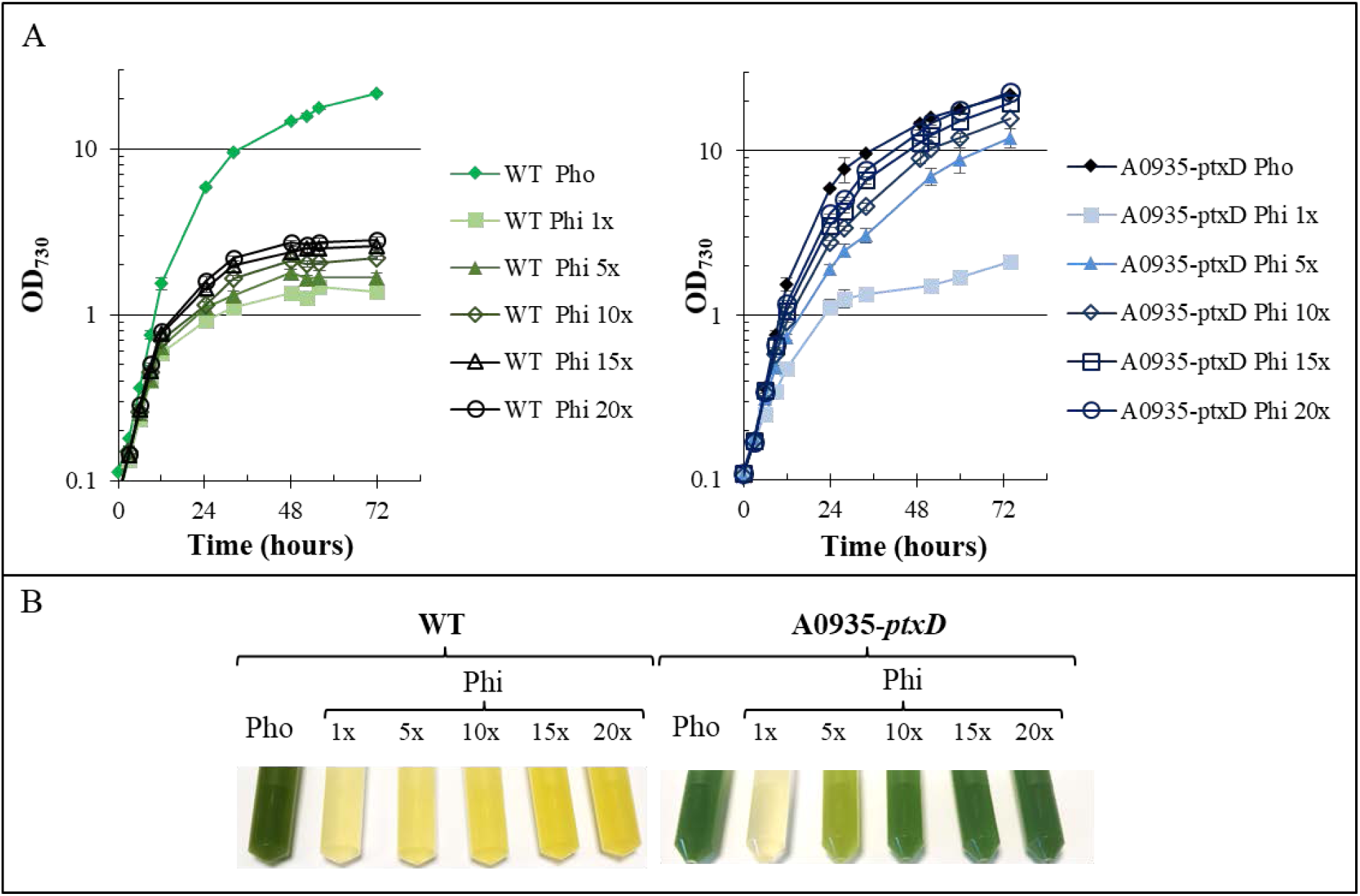
Growth of phosphite utilizing strains in phosphate (Pho) and phosphite (Phi) containing AD7 medium **A.** Growth curve of WT Syn7002 (left) and phosphite (right) utilizing strain in AD7 medium containing Pho or different concentrations of Phi, as indicated. **B.** Detail of culture samples at 48 hours after inoculation. Factors for converting OD_730_ to grams dry cell weight (gDCW)·L^−1^ can be found in Suppl. Table 2.

Finally, we mutated the RBS sequence upstream of the *triA* gene in the original pSJ051 to match the mutation found in Mel5. Using this modified construct we were able to obtain colonies by plating directly on AD7-Mel plates (as well as by plating on AD7-Cya plates). Growth of the newly obtained strain (named “Re-Mel5”) was compared to Mel5 and Re-Mel5 was found to reach a similar OD_730_ after 48 hours when grown on melamine as the sole nitrogen source (Suppl. Fig. 4), indicating that the mutation found was indeed able to improve melamine utilization efficiency.

### 3.2. Phosphite and PtxD can be used as an efficient selection system in Synechococcus sp. PCC7002

Though phosphite (Phi) was previously shown to be able to sustain growth of modified strains of both *Synechocystis* sp. PCC 6803 (Polyviou et al., 2015) and *Synechococcus* sp. PCC 7942 (Motomura et al., 2018), in both cases the genetic manipulations (integration of the operon containing both *ptxD* as well as a specific Phi transporter) were driven by antibiotic selection pressure. At the same time none of the two strains (neither wild-type *Synechocystis* sp. PCC 6803 nor wild-type *Synechococcus* sp. PCC 7942) seem to be able to take up Phi without inclusion of a specific transporter, thus making the construct too large to be of practical use as a selectable marker. As no data exist in the literature regarding the ability of Syn7002 to grow on Phi as sole P source, we tested growth of the WT strain using varying concentrations of Phi (Fig. 3A, left panel). Though there was some growth for the first 24-36 hours, no observable growth occurred after this period, which was probably due to the full consumption of internal phosphate reserves – cyanobacteria store phosphorus as polyphosphate granules in the cytoplasm and have a dynamic mobilization mechanism that allows them to tap onto these reserves when needed (Gomez-Garcia et al., 2013).

Syn7002 WT transformed with a construct (pSJ135) containing the *Pseudomonas stutzeri* WM88 phosphite dehydrogenase gene in the neutral site *A0935* and no other selectable marker (Fig. 4A, top) was plated on AD7 plates with Phi (0.37 mM, 1x) as sole phosphorus source. This transformation yielded many hundred colonies, and, though the transformants (A0935-ptxD) had a yellowish tinge, characteristic of phosphate-starved cells, this concentration of Phi was sufficient to induce full chromosomal segregation of the transformed strains (Fig. 4B), thus validating the *ptxD* gene and phosphite as a valid selection strategy in Syn7002.

Given that the *ptxD* gene alone was enough to permit growth of strain A0935-ptxD on Phi, it would seem that, unlike the related (freshwater) strains *Synechocystis* sp. PCC6803 and *Synechococcus* sp. PCC7942, Syn7002 is able to import Phi from the growth medium, through a so far undefined transporter.

To test whether a higher concentration gradient would be sufficient to enhance transport into the cells and allow faster growth on Phi, growth of A0935-ptxD was tested in AD7 with increasing Phi concentrations (from 0.37 mM to 7.4 mM). As can be seen in Fig. 3A (right side) the highest Phi concentration tested (Phi 20x, 7.4 mM) allowed this strain to attain growth rates nearing those of Pho-grown cells. At the same time, the relation between grams dry cell weight (gDCW) and OD_730_ also increased with increasing Phi concentrations, from 0.1633±0.014 gDCW·L^−1^·OD_730_^−1^ for cells grown in AD7-Phi 1x to 0.2038±0.003 gDCW·L^−1^·OD_730_^−1^ for cells grown in AD7-Phi 20x (Suppl. Table 2), a value similar to that of WT cells grown in standard conditions (0.2145±0.007 gDCW·L^−1^·OD_730_^−1^). This increase might be due to the more efficient Phi uptake and conversion at higher Phi concentrations.

Previous work demonstrated that transgenic *Arabidopsis thaliana* plants expressing the same *ptxD* gene (using phosphinothricin for selection) were also able to grow using Phi as the sole phosphorus source (Lopez-Arredondo and Herrera-Estrella, 2012). While the specific transporter by which Phi is taken up by plant roots is not known, it would seem that no extra Phi transporter genes are required, as is the case for Syn7002. However, Phi uptake in Syn7002, unlike *A. thaliana*, is much less efficient. As previous studies have shown that several marine cyanobacteria are able to take up and utilize Phi as a phosphorus source (Feingersch et al., 2012; Martinez et al., 2012; Polyviou et al., 2015) we searched the Syn7002 genome for putative transporter genes, such as *ptxB* and *phnD* homologues (Bisson et al., 2017). While no *ptxB* homologues could be found, two putative *phnD* genes, *A0336* and *G0143*, were present in either the circular chromosome (*A0336*) or the pAQ7 plasmid (*G0143*) (see Suppl. Fig. 5). We hypothesized that either of these two genes might be involved in phosphite import to the cell, as *phnD* homologues were shown to also bind phosphite (Bisson et al., 2017). However, knock-out of these putative phosphonate transporters, either alone or in combination, did not result in an inability of the *ptxD* parental strain to grow on phosphite (Suppl. Fig. 6). As such, further studies will be required in order to more fully understand phosphite transport by Syn7002.

**Figure 4.**
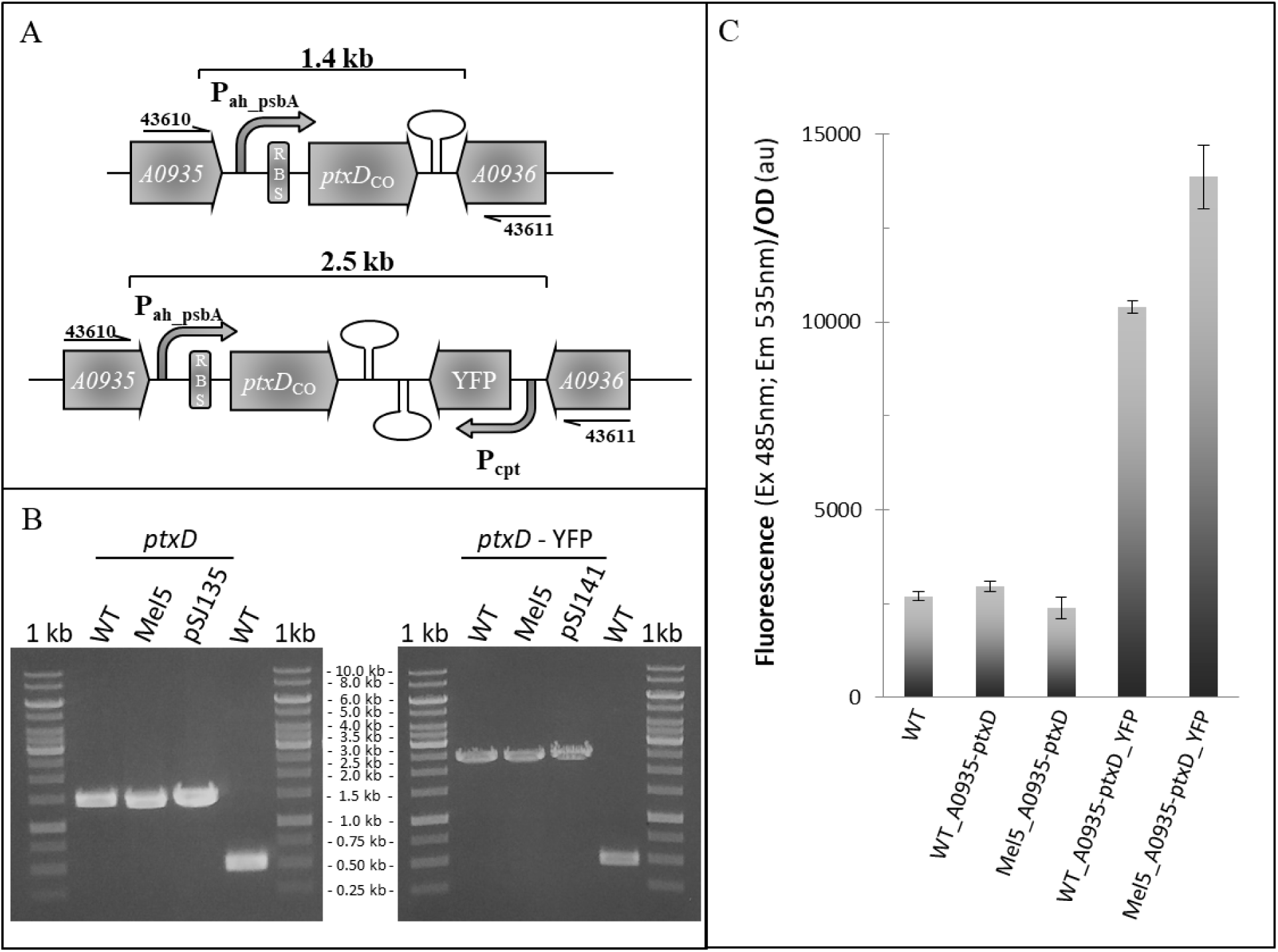
Overview of the phosphite selection tool. **A.** *Top* -Detail of pSJ135, including primers using for chromosomal integration PCR. *Bottom* – Detail of construct pSJ141, which uses phosphite to drive chromosomal integration of a heterologous gene (YFP). **B.** 0.8% agarose gel of PCR showing genome integration of ptxD gene and ptxD-driven integration of the YFP gene, in both WT and Mel5 backgrounds. **C.** YFP fluorescence in strains transformed with pSJ135 and pSJ141 vs. respective background strains.

**Figure 5.**
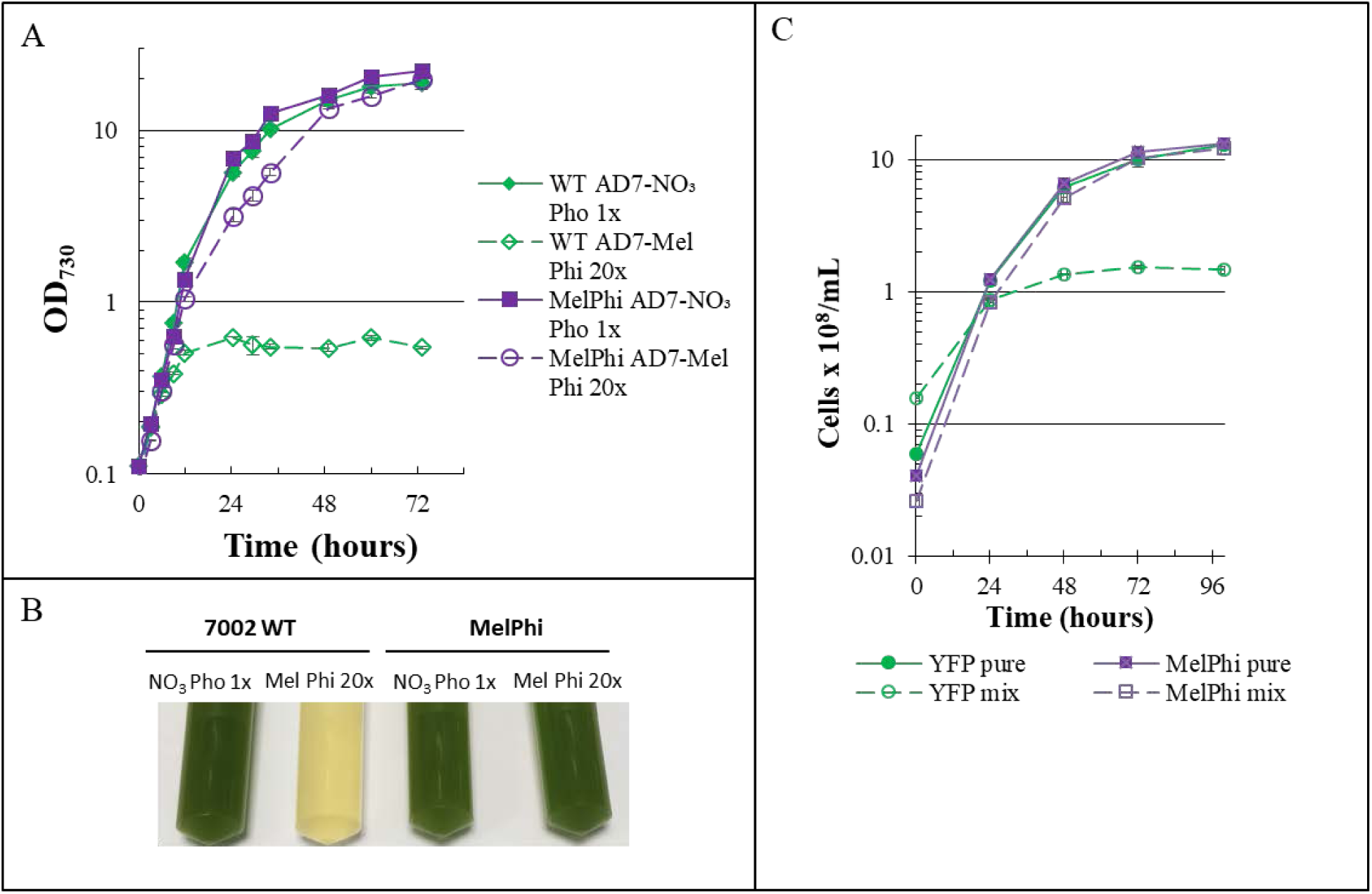
Characterization of the melamine and phosphite utilizing strain. **A.** Growth curves of WT and Mel5-A0935ptxD (“MelPhi”) strains in either regular AD7 medium or AD7-Mel Phi 20x. **B.** Detail of culture samples at 48 hours post inoculation. Factors for OD_730_ to grams dry cell weight (gDCW)·L^−1^ conversion were calculated for the MelPhi strain in AD7-Mel Pho 1x or AD7-Mel Phi 20x and can be found in Suppl. Table 2. **C.** Growth curves of a strain expressing YFP in the Syn7002 WT background (“YFP pure”, grown in regular AD7), the MelPhi strain (“MelPhi pure”, lacking YFP, grown in AD7-Mel Phi 20x), or mixed cultures of the two strains (“YFP mix” and “MelPhi mix”), combined in at a cell ratio of 6:1 YFP (in WT background) to MelPhi (lacking YFP), in AD7-Mel Phi 20x, measured by flow cytometry.

To investigate whether this selection method would be sufficient to allow co-integration of other genes, we designed a second construct where the *yfp* gene would be integrated into the chromosome at the same locus, using *ptxD* as the selectable marker and Phi as positive selection. As can be seen in Figs. 4B and 4C, the *yfp* gene was successfully co-integrated using this method and YFP fluorescence could be measured in positive transformants. Thus, this method can be used to select for integration and expression of heterologous genes in Syn7002.

### 3.3. Construction of a strain capable of growing on both melamine and phosphite

As we had now validated two separate antibiotic-free selection methods in Syn7002, we hypothesized that a double selection, using both melamine and Phi, should be possible in the same chassis. Therefore we proceeded to transform Mel5, one of the best melamine-growing strains, with either pSJ135 (*ptxD* gene alone) or pSJ141 (*ptxD* and *yfp*) and selecting for positive transformants on AD7-Mel 1xPhi plates. As was the case for the Syn7002 WT background, positive transformants could be readily obtained by this selection method (Fig. 4B) and YFP fluorescence measured in the fully segregated YFP-expressing transformants (Fig. 4C). Strain Mel5-A0935ptxD (“MelPhi”) was able to grow in AD7-Mel Phi 20x liquid medium, albeit at a slightly slower rate, in comparison to MelPhi or the Syn7002 WT parental strain grown in regular AD7 (Fig. 5A). As predicted, Syn7002 WT was unable to grow in AD7-Mel Phi 20x medium (Fig. 5A).

### 3.4. Strain Mel5-A0935ptxD is able to resist deliberate contamination

As mentioned earlier, the overarching aim of this work was to develop a cyanobacterial strain that would be suitable for outdoor cultivation in open systems. This strain should, therefore, be able to outcompete other strains under these conditions, so as to become the dominant population in a potentially contaminated system. To determine the robustness of the obtained strain, an experiment was devised where the starting culture for Mel5-A0935ptxD (in AD7-Mel Phi 20x) was deliberately contaminated with a large excess, either 6 times (Fig. 5C, top) or 10 times higher cell counts (Suppl. Fig 5) of a Syn7002 strain constitutively expressing YFP (from the P_cpt_ promoter). As can be seen in Fig. 5C, Mel5-A0935ptxD (“MelPhi”) was able to overcome this large excess of contaminant and become the dominant population, thus showing that this strain is indeed suitable for outdoor cultivation under unsterilized conditions. Though, as discussed above, there is a leakage of melamine pathway intermediates from the parental strain Mel5 cells during cultivation in melamine (Suppl. Fig. 3), the low amounts of these xenobiotic compounds released by the cells (at maximum 0.2 mM) are very unlikely to support bacterial growth, thus negating any form of scavenging from contaminant cells. The slight growth of the YFP contaminant observed is, as mentioned above (see section 3.2), most likely due to mobilization and consumption of internal nutrient reserves, though this is unable to sustain long term growth. Previously, transgenic *A. thaliana* expressing *ptxD* was also shown to be able to resist and overcome deliberate contamination by common weeds (Lopez-Arredondo and Herrera-Estrella, 2012), underscoring the advantage of this approach to give strains grown in unsterilized conditions an edge over the competition.

One important consideration of the strains developed in this study is that, at current prices and at the concentrations utilized in this study, melamine would be a more economical nitrogen source than nitrate, with a 24% reduction in cost when using melamine (see Suppl. Table 3). Also, as melamine is not used as an agricultural fertilizer, its usage as nitrogen source would eliminate competition for nitrogen-rich fertilizers used in agriculture. Furthermore, as melamine levels drop to below the level of detection using LC-MS/MS within 24 hours of growth, residual melamine in the final culture supernatant would not be a deterring factor in adoption of this technique.

The additional selection by phosphite gives the strain a “double edge” that will be even more difficult to overcome by contaminating species, especially in the early stages of the cultures, thus allowing strains carrying these two modifications to become the dominant population without the need for sterilization or antibiotic addition. Furthermore, this strategy negates the risk for horizontal gene transfer of antibiotic resistance cassettes, a growing problem in today’s society (Ventola, 2015a; Ventola, 2015b; von Wintersdorff et al., 2016). Though there is still the risk of horizontal gene transfer of the introduced pathways into environmental microorganisms, these pathways would be very unlikely to become a risk to either the ecological balance of local microorganism populations or human health. However, to completely eliminate such a risk, a biocontainment strategy for both the melamine operon as well as the Phi strategy, similar to that described for Phi usage in *E. coli* (Hirota et al., 2017) and *Synechococcus* sp. PCC 7942 (Motomura et al., 2018) can also be implemented by deleting the cognate ammonium, nitrate and phosphate transporter genes.

At the current stage using phosphite in large cultures would be uneconomical (see Suppl. Table 3), due to the need for a large concentration excess being required for efficient growth. However, this could be mitigated by introducing a more efficient transport system, such as the one previously described for *S. elongatus* sp. PCC 7942 (Motomura et al., 2018) or using phosphite as an initial selection marker, before switching to phosphate for large scale growth.

In conclusion, this work describes, for the first time, marine cyanobacterial strains that are able to grow on melamine as sole nitrogen source and the use of phosphite selection as an efficient selection strategy in cyanobacteria. Finally, we developed a unique strain that is able to use both melamine and phosphite as sole N and P sources. This strain is able to resist deliberate contamination by other cyanobacteria, even when the contamination is present in large excess, and should prove to be a useful chassis strain for “green” biotechnological applications.

## Supporting information

Supplemental Table 1 - primers

Supplemental Table 2 - OD to gDCW

Supplemental Table 3 - price for N and P sources

## Abbreviations

WT: Wild type
YFP: Yellow Fluorescent Protein
RBS: Ribosome Binding Site

## Acknowledgements

This work was supported by NTU grants M4080306 to BN and M4081714 to PJN. The authors would like to thank Daniela Moses (SCELCE, NTU) for assistance with whole genome sequencing reactions and the NTU Phenomics Centre for LC-MS/MS analysis of culture supernatants.

## References

Angermayr, S. A., Paszota, M., Hellingwerf, K. J., 2012. Engineering a cyanobacterial cell factory for production of lactic acid. Applied and environmental microbiology. 78, 7098–106.10.1128/AEM.01587-12

Arnon, D. I., McSwain, B. D., Tsujimoto, H. Y., Wada, K., 1974. Photochemical activity and components of membrane preparations from blue-green algae. I. Coexistence of two photosystems in relation to chlorophyll a and removal of phycocyanin. Biochim Biophys Acta. 357, 231–45

Begemann, M. B., Zess, E. K., Walters, E. M., Schmitt, E. F., Markley, A. L., Pfleger, B. F., 2013. An organic acid based counter selection system for cyanobacteria. PLoS One. 8, e76594. 10.1371/journal.pone.0076594

Bisson, C., Adams, N. B. P., Stevenson, B., Brindley, A. A., Polyviou, D., Bibby, T. S., Baker, P. J., Hunter, C. N., Hitchcock, A., 2017. The molecular basis of phosphite and hypophosphite recognition by ABC-transporters. Nat Commun. 8, 1746.10.1038/s41467-017-01226-8

Braekevelt, E., Lau, B. P., Feng, S., Menard, C., Tittlemier, S. A., 2011. Determination of melamine, ammeline, ammelide and cyanuric acid in infant formula purchased in Canada by liquid chromatography-tandem mass spectrometry. Food Addit Contam Part A Chem Anal Control Expo Risk Assess. 28, 698–704.10.1080/19440049.2010.545442

Choi, S. Y., Wang, J. Y., Kwak, H. S., Lee, S. M., Um, Y., Kim, Y., Sim, S. J., Choi, J. I., Woo, H. M., 2017. Improvement of squalene production from CO_2_ in *Synechococcus elongatus* PCC 7942 by metabolic engineering and scalable production in a photobioreactor. ACS Synth Biol. 6, 1289–1295.10.1021/acssynbio.7b00083

Clark, R. L., McGinley, L. L., Purdy, H. M., Korosh, T. C., Reed, J. L., Root, T. W., Pfleger, B. F., 2018. Light-optimized growth of cyanobacterial cultures: Growth phases and productivity of biomass and secreted molecules in light-limited batch growth. Metab Eng. 47, 230–242.10.1016/j.ymben.2018.03.017

Collier, J. L., Grossman, A. R., 1992. Chlorosis induced by nutrient deprivation in *Synechococcus* sp. strain PCC 7942: not all bleaching is the same. Journal of bacteriology. 174, 4718–26

Davies, F. K., Work, V. H., Beliaev, A. S., Posewitz, M. C., 2014. Engineering limonene and bisabolene production in wild type and a glycogen-deficient mutant of *Synechococcus* sp. PCC 7002. Frontiers in bioengineering and biotechnology. 2, 21.10.3389/fbioe.2014.00021

Dexter, J., Armshaw, P., Sheahan, C., Pembroke, J. T., 2015. The state of autotrophic ethanol production in Cyanobacteria. J Appl Microbiol. 119, 11–24.10.1111/jam.12821

Elhai, J., Wolk, C. P., 1988. A versatile class of positive-selection vectors based on the nonviability of palindrome-containing plasmids that allows cloning into long polylinkers. Gene. 68, 119–138.Doi10.1016/0378-1119(88)90605-1

Englund, E., Pattanaik, B., Ubhayasekera, S. J., Stensjo, K., Bergquist, J., Lindberg, P., 2014. Production of squalene in *Synechocystis* sp. PCC 6803. PLoS One. 9, e90270.10.1371/journal.pone.0090270

Fathima, A. M., Chuang, D., Lavina, W. A., Liao, J., Putri, S. P., Fukusaki, E., 2018. Iterative cycle of widely targeted metabolic profiling for the improvement of 1-butanol titer and productivity in *Synechococcus elongatus*. Biotechnol Biofuels. 11, 188.10.1186/s13068-018-1187-8

Feingersch, R., Philosof, A., Mejuch, T., Glaser, F., Alalouf, O., Shoham, Y., Beja, O., 2012. Potential for phosphite and phosphonate utilization by *Prochlorococcus*. ISME J. 6, 827–34. 10.1038/ismej.2011.149

Frigaard, N. U., Sakuragi, Y., Bryant, D. A., 2004. Gene inactivation in the cyanobacterium *Synechococcus* sp. PCC 7002 and the green sulfur bacterium *Chlorobium tepidum* using in vitro-made DNA constructs and natural transformation. Methods Mol Biol. 274, 325–40. 10.1385/1-59259-799-8:325

Gomez-Garcia, M. R., Fazeli, F., Grote, A., Grossman, A. R., Bhaya, D., 2013. Role of polyphosphate in thermophilic *Synechococcus* sp. from microbial mats. Journal of bacteriology. 195, 3309–19.10.1128/JB.00207-13

Gordon, G. C., Korosh, T. C., Cameron, J. C., Markley, A. L., Begemann, M. B., Pfleger, B. F., 2016. CRISPR interference as a titratable, trans-acting regulatory tool for metabolic engineering in the cyanobacterium *Synechococcus* sp. strain PCC 7002. Metab Eng. 38, 170–179.10.1016/j.ymben.2016.07.007

Halfmann, C., Gu, L., Gibbons, W., Zhou, R., 2014. Genetically engineering cyanobacteria to convert CO_2_, water, and light into the long-chain hydrocarbon farnesene. Appl Microbiol Biotechnol. 98, 9869–77.10.1007/s00253-014-6118-4

Hirota, R., Abe, K., Katsuura, Z. I., Noguchi, R., Moribe, S., Motomura, K., Ishida, T., Alexandrov, M., Funabashi, H., Ikeda, T., Kuroda, A., 2017. A novel biocontainment strategy makes bacterial growth and survival dependent on phosphite. Sci Rep. 7, 44748.10.1038/srep44748

Kanda, K., Ishida, T., Hirota, R., Ono, S., Motomura, K., Ikeda, T., Kitamura, K., Kuroda, A., 2014. Application of a phosphite dehydrogenase gene as a novel dominant selection marker for yeasts. J Biotechnol. 182, 68–73.10.1016/j.jbiotec.2014.04.012

Kato, A., Takatani, N., Ikeda, K., Maeda, S. I., Omata, T., 2017. Removal of the product from the culture medium strongly enhances free fatty acid production by genetically engineered *Synechococcus elongatus*. Biotechnol Biofuels. 10, 141.10.1186/s13068-017-0831-z

Liu, H., Naismith, J. H., 2008. An efficient one-step site-directed deletion, insertion, single and multiple-site plasmid mutagenesis protocol. BMC Biotechnol. 8, 91.10.1186/1472-6750-8-91

Loera-Quezada, M. M., Leyva-Gonzalez, M. A., Velazquez-Juarez, G., Sanchez-Calderon, L., Do Nascimento, M., Lopez-Arredondo, D., Herrera-Estrella, L., 2016. A novel genetic engineering platform for the effective management of biological contaminants for the production of microalgae. Plant Biotechnol J. 14, 2066–76.10.1111/pbi.12564

Lopez-Arredondo, D. L., Herrera-Estrella, L., 2012. Engineering phosphorus metabolism in plants to produce a dual fertilization and weed control system. Nat Biotechnol. 30, 889–93.10.1038/nbt.2346

Ludwig, M., Bryant, D. A., 2011. Transcription profiling of the model cyanobacterium *Synechococcus* sp. strain PCC 7002 by next-gen (SOLiD) sequencing of cDNA. Frontiers in microbiology. 2, 41

Ludwig, M., Bryant, D. A., 2012. Synechococcus sp. strain PCC 7002 transcriptome: acclimation to temperature, salinity, oxidative stress, and mixotrophic growth conditions. Frontiers in microbiology. 3, 354

Markley, A. L., Begemann, M. B., Clarke, R. E., Gordon, G. C., Pfleger, B. F., 2015. Synthetic biology toolbox for controlling gene expression in the cyanobacterium Synechococcus sp. strain PCC 7002. ACS Synth Biol. 4, 595–603.10.1021/sb500260k

Martinez, A., Osburne, M. S., Sharma, A. K., DeLong, E. F., Chisholm, S. W., 2012. Phosphite utilization by the marine picocyanobacterium *Prochlorococcus* MIT9301. Environ Microbiol. 14, 1363–77.10.1111/j.1462-2920.2011.02612.x

Motomura, K., Sano, K., Watanabe, S., Kanbara, A., Gamal Nasser, A. H., Ikeda, T., Ishida, T., Funabashi, H., Kuroda, A., Hirota, R., 2018. Synthetic phosphorus metabolic pathway for biosafety and contamination management of cyanobacterial cultivation. ACS Synth Biol. 10.1021/acssynbio.8b00199

Nahampun, H. N., Lopez-Arredondo, D., Xu, X., Herrera-Estrella, L., Wang, K., 2016. Assessment of *ptxD* gene as an alternative selectable marker for *Agrobacterium*-mediated maize transformation. Plant Cell Rep. 35, 1121–1132.10.1007/s00299-016-1942-x

Pandeya, D., Campbell, L. M., Nunes, E., Lopez-Arredondo, D. L., Janga, M. R., Herrera-Estrella, L., Rathore, K. S., 2017. *ptxD* gene in combination with phosphite serves as a highly effective selection system to generate transgenic cotton (*Gossypium hirsutum* L.). Plant Mol Biol. 95, 567–577. 10.1007/s11103-017-0670-0

Perez, A. A., Liu, Z., Rodionov, D. A., Li, Z., Bryant, D. A., 2016. Complementation of cobalamin auxotrophy in *Synechococcus* sp. strain PCC 7002 and validation of a putative cobalamin riboswitch in vivo. Journal of bacteriology.10.1128/JB.00475-16

Polyviou, D., Hitchcock, A., Baylay, A. J., Moore, C. M., Bibby, T. S., 2015. Phosphite utilization by the globally important marine diazotroph *Trichodesmium*. Environmental microbiology reports. 7, 824–30.10.1111/1758-2229.12308

Ruffing, A. M., 2014. Improved free fatty acid production in cyanobacteria with *Synechococcus* sp. PCC 7002 as host. Frontiers in bioengineering and biotechnology. 2, 17.10.3389/fbioe.2014.00017

Schoepp, N. G., Stewart, R. L., Sun, V., Quigley, A. J., Mendola, D., Mayfield, S. P., Burkart, M. D., 2014. System and method for research-scale outdoor production of microalgae and cyanobacteria. Bioresour Technol. 166, 273–81.10.1016/j.biortech.2014.05.046

Shabestary, K., Anfelt, J., Ljungqvist, E., Jahn, M., Yao, L., Hudson, E. P., 2018. Targeted repression of essential genes to arrest growth and increase carbon partitioning and biofuel titers in cyanobacteria. ACS Synth Biol. 7, 1669–1675.10.1021/acssynbio.8b00056

Shaw, A. J., Lam, F. H., Hamilton, M., Consiglio, A., MacEwen, K., Brevnova, E. E., Greenhagen, E., LaTouf, W. G., South, C. R., van Dijken, H., Stephanopoulos, G., 2016. Metabolic engineering of microbial competitive advantage for industrial fermentation processes. Science. 353, 583–6. 10.1126/science.aaf6159

Stevens, S. E., Patterson, C. O., Myers, J., 1973. Production of hydrogen peroxide by blue-green algae - a survey. J Phycol. 9, 427–430

Ventola, C. L., 2015a. The antibiotic resistance crisis: part 1: causes and threats. P T. 40, 277–83

Ventola, C. L., 2015b. The antibiotic resistance crisis: part 2: management strategies and new agents. P T. 40, 344–52

von Wintersdorff, C. J. H., Penders, J., van Niekerk, J. M., Mills, N. D., Majumder, S., van Alphen, L. B., Savelkoul, P. H. M., Wolffs, P. F. G., 2016. Dissemination of antimicrobial resistance in microbial ecosystems through horizontal gene transfer. Frontiers in microbiology. 7.10.3389/fmicb.2016.00173

Wang, X., Liu, W., Xin, C., Zheng, Y., Cheng, Y., Sun, S., Li, R., Zhu, X. G., Dai, S. Y., Rentzepis, P. M., Yuan, J. S., 2016. Enhanced limonene production in cyanobacteria reveals photosynthesis limitations. Proceedings of the National Academy of Sciences of the United States of America. 113, 14225–14230.10.1073/pnas.1613340113

Xu, Y., Alvey, R. M., Byrne, P. O., Graham, J. E., Shen, G., Bryant, D. A., 2011. Expression of genes in cyanobacteria: adaptation of endogenous plasmids as platforms for high-level gene expression in Synechococcus sp. PCC 7002. Methods Mol Biol. 684, 273–93

