## Supplemental Figures for "Growth and selection of the cyanobacterium *Synechococcus* sp. PCC 7002 using alternative nitrogen and phosphorus sources"

Supplementary figures

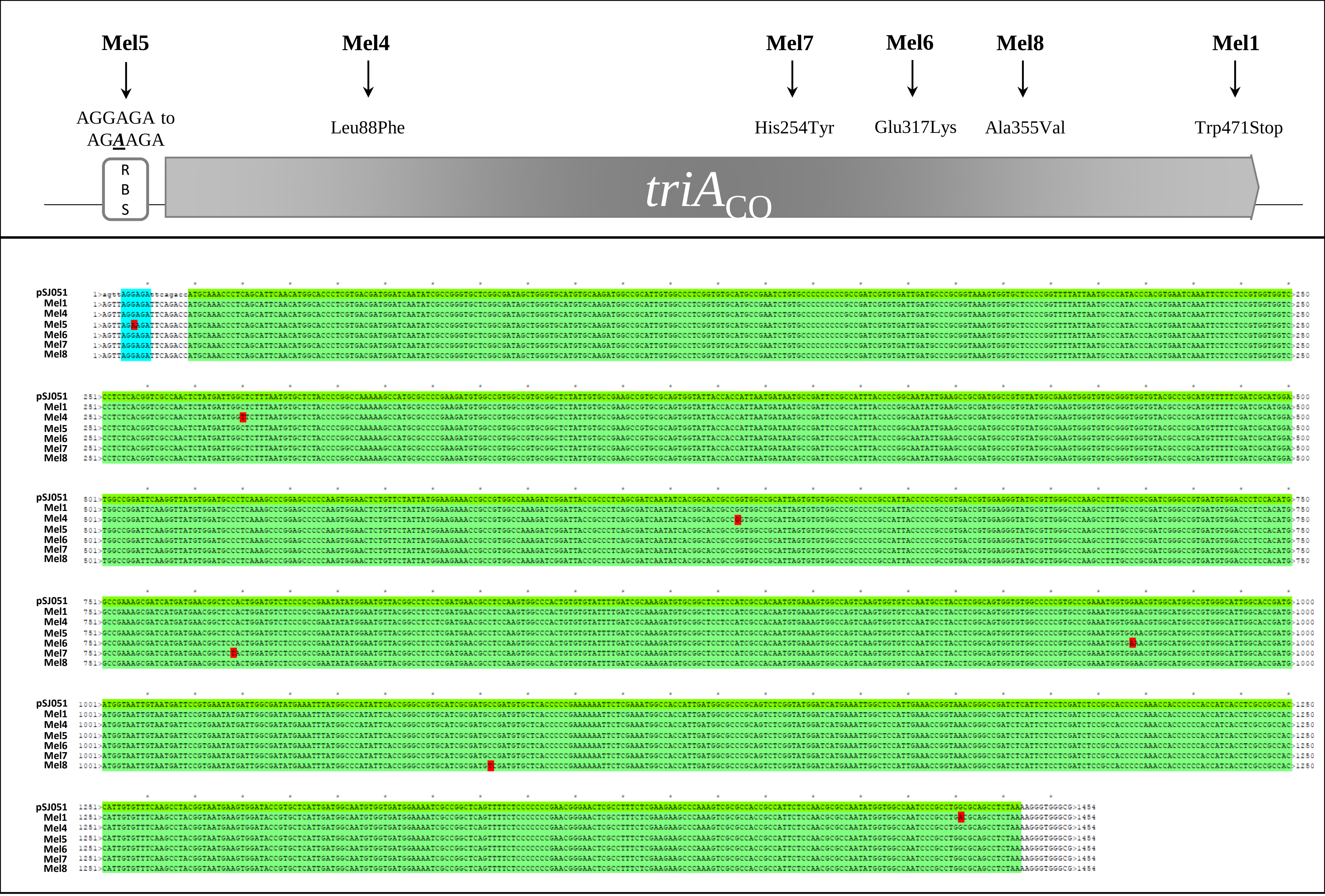


**Supp. Fig. 1** – *Top* - Schematic representation of mutations to the *triA* locus in different melamine utilizing strains, as found by Illumina sequencing. Mel1 has a mutation 4 amino acids before the original stop codon. *Bottom* – Multiple sequence alignment of the *triA* locus in the different melamine utilizing strains

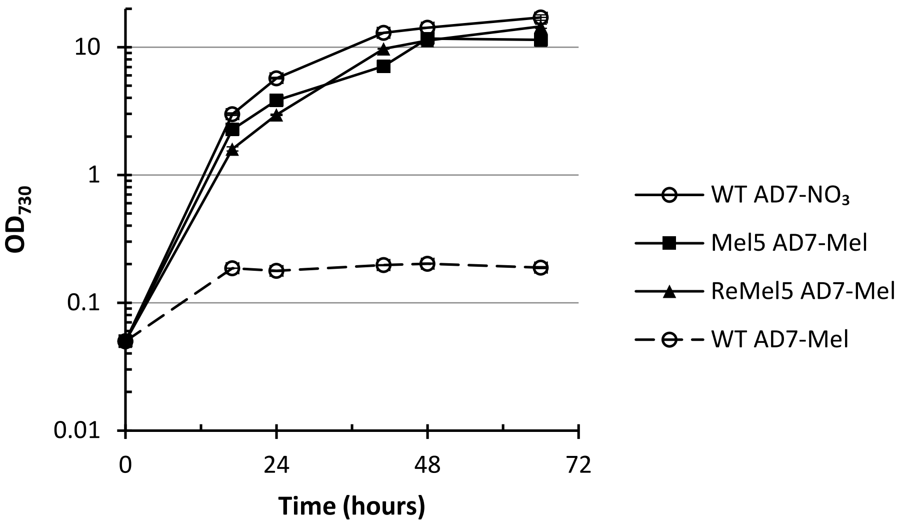


**Suppl. Fig. 2** – Growth of WT, Mel5 and Re-Mel5 strains in either normal AD7-$NO_3$ or AD7-Mel

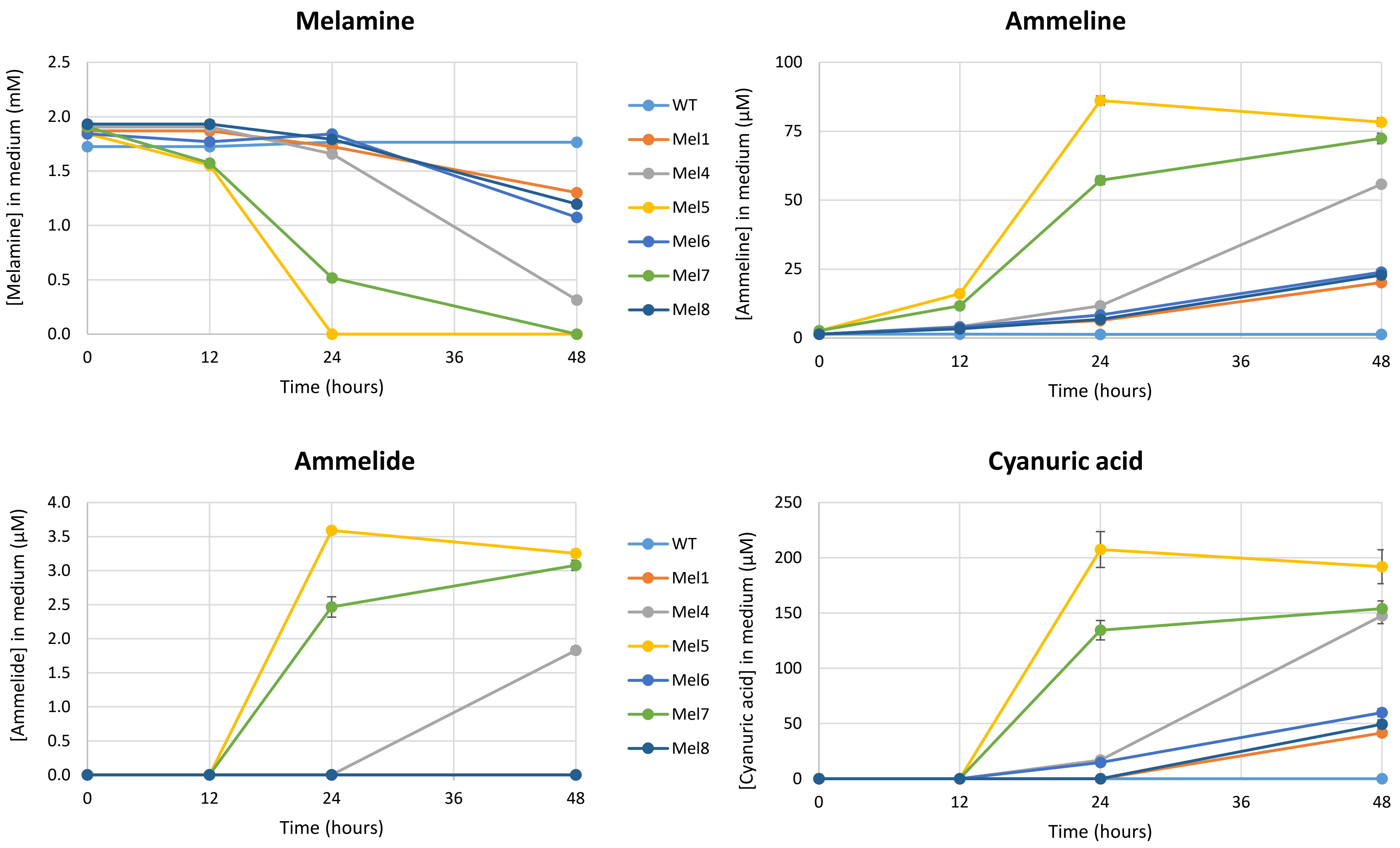


**Suppl. Fig. 3** – LC-MS/MS quantification of melamine pathway intermediates in spent culture medium, at the time points indicated. Quantification for WT culture inoculated in AD7-Mel medium is also included, as a control. Notice the difference in scale for melamine (in mM) and remaining intermediates (in µM). Error bars may not be apparent due to scale.

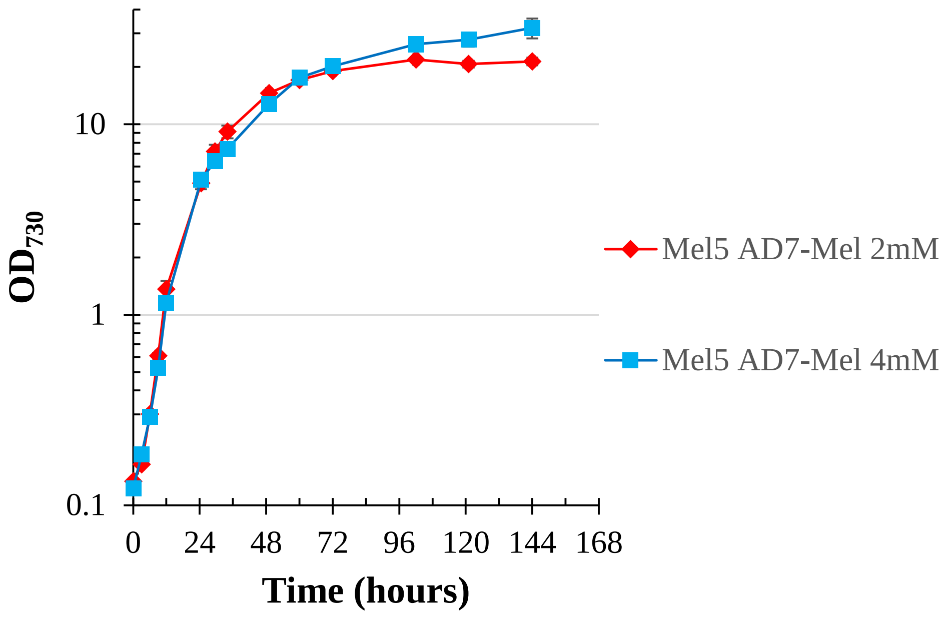


**Suppl. Fig. 4** – Growth curves for the Mel5 strain, grown in AD7-Mel medium containing either 2 mM or 4 mM melamine

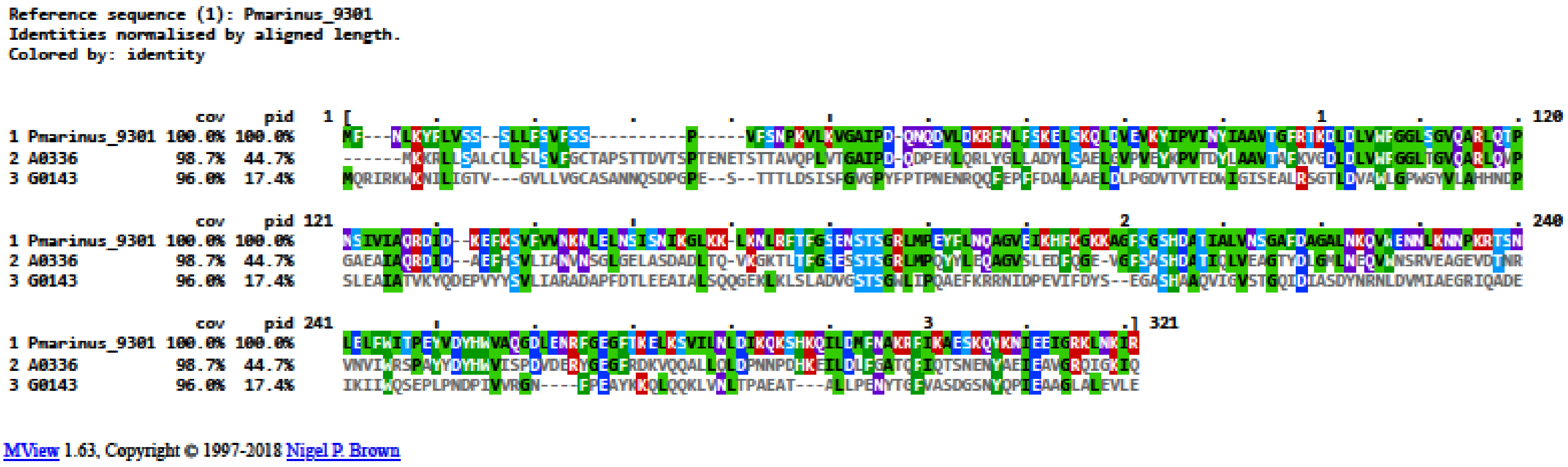


**Suppl. Fig. 5** – ClustalOmega alignment of PhnD from *Prochlorococcus marinus* MIT9301 and the two putative homologues in Syn7002 (visualized and coloured using MView)

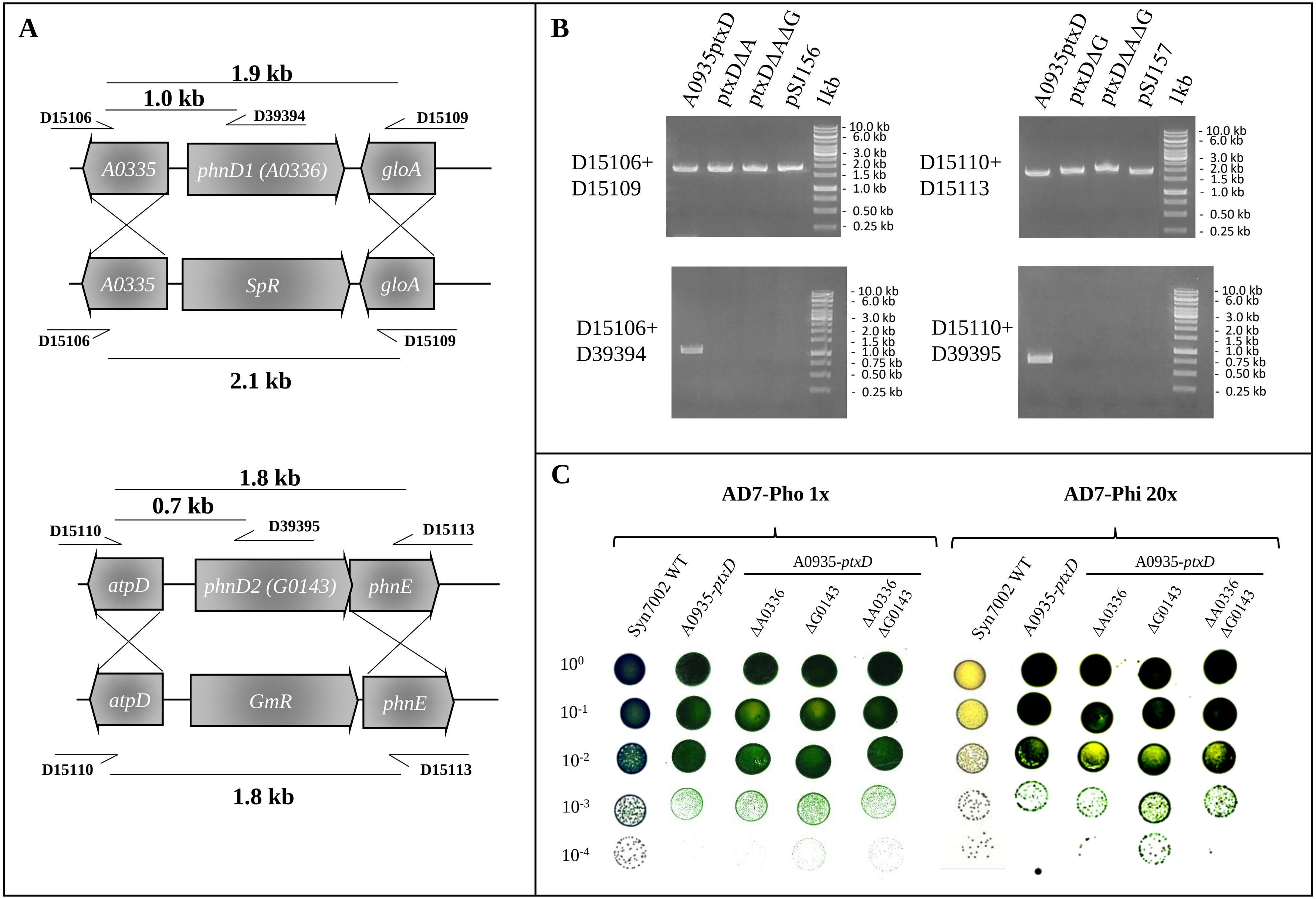


**Suppl. Fig. 6** – Knock-out of putative phosphonate transporter homologues *A0336* (top) and *G0143* (bottom) in Syn7002. **A.** Schematic representation of knock-out construct plasmids pSJ156 (top) and pSJ157 (bottom). Individual elements are not to scale. **B.** Segregation gels for A0935-ptxD putative phosphonate transporter homologue knock-out strains. **C.** Dilution plating of A0935-ptxD parental strain and derivative knock-out strains, in either AD7-Pho 1x (*left*) or AD7-Phi 20x (*right*).
Note: ΔA – Δ*A0336*::*SpR;* ΔG – Δ*G0143*::*GmR*

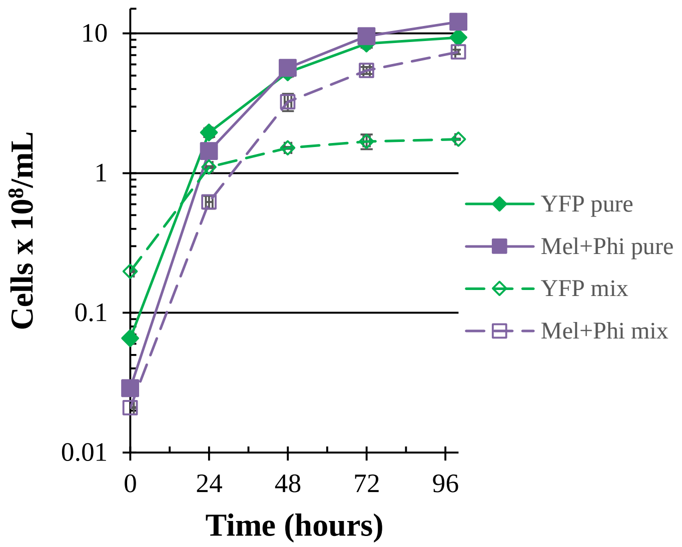


**Suppl. Fig. 7** – Growth curves of a strain expressing YFP in the Syn7002 WT background (“YFP pure”, grown in regular AD7), the MelPhi strain (“MelPhi pure”, lacking YFP, grown in AD7-Mel Phi 20x), or mixed cultures of the two strains (“YFP mix” and “MelPhi mix”), combined in at a cell ratio of 10:1 YFP (in WT background) to MelPhi (lacking YFP), in AD7-Mel Phi 20x, measured by flow cytometry.

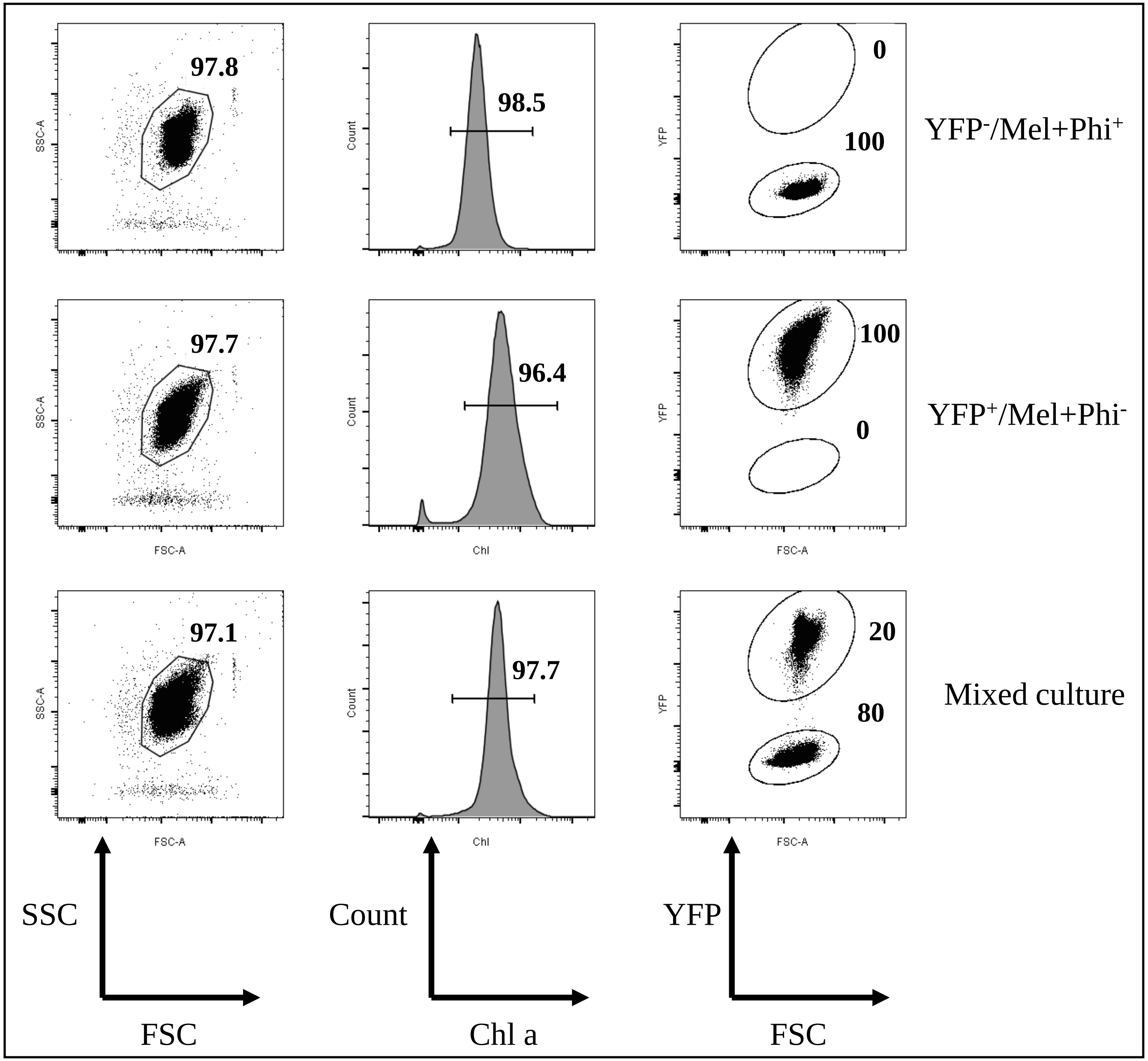


**Suppl. Fig. 8** – Gating strategy used for cell counts in the contamination experiment. Data shown is for one of the 10:1 mixed culture experiments, at T=100 hours. *Left* – Gates drawn on dot plots for (top to bottom) YFP and Mel+Phi pure cultures and the mixed culture; *Middle* – Histogram plots for the same samples; *Right* – Dot plots for the same samples using YFP vs. forward scatter, used for quantification. SSC – side scatter; FSC – forward scatter

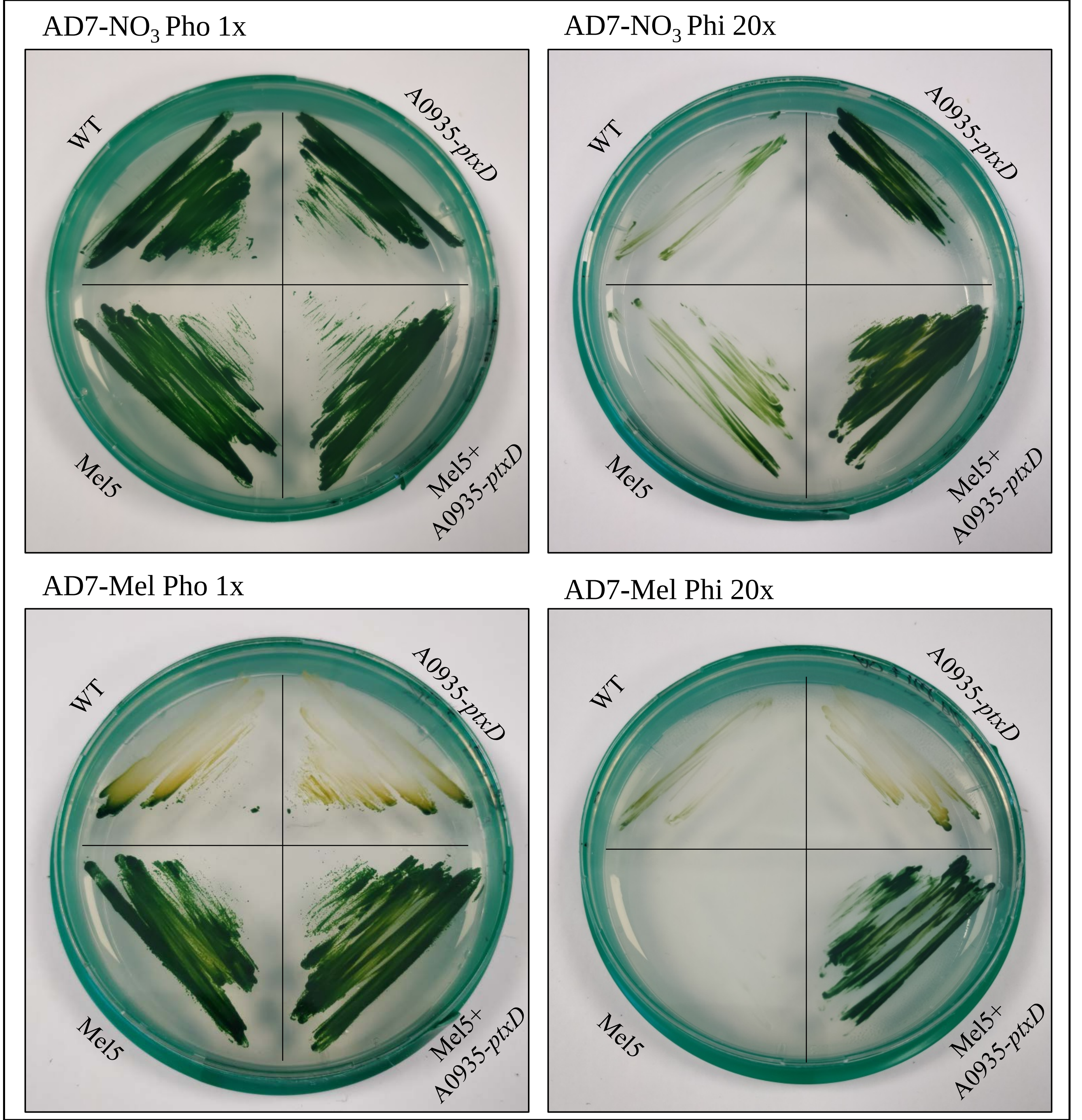


**Suppl. Fig. 9** – Growth of Syn7002 WT and melamine and phosphite utilizing strains in AD7 with either nitrate ($NO_3$) or melamine (Mel) and phosphate (Pho) or phosphite (Phi). All plates were grown at 30 °C, 80 $\mu E \cdot m^{-2} \cdot s^{-1}$ and 1% $CO_2$ for 5 days
